# LIT (Layer-Wise Image Trajectories): In Situ Monitoring for Early Quality Prediction and Anomaly Detection in Acellular and Cell-Laden Two-Photon Polymerization

**DOI:** 10.64898/2026.08.14.744878

**Authors:** Egon Prioglio, Chiara Scrocciolani, Bianca Maria Colosimo

## Abstract

Two-photon polymerization (2PP) enables fabrication of hydrogel constructs with submicron, cell-scale resolution, but hydrogel-based bioinks are markedly more sensitive to process variability than conventional photoresists, and this sensitivity is further amplified when living cells are embedded in the resin. Post-processing evaluation, performed only after development, occurs too late to enable any corrective action. A full-factorial design of experiments across laser power and scan speed shows that fabrication outcome depends on both parameter choice and cell presence, with cells shifting and broadening the range of conditions yielding structurally sound constructs. However, substantial variability persists within each nominal condition and cannot be resolved by parameter refinement alone, indicating that outcome is governed by what occurs during each individual print rather than by the parameters set. To capture this, a layer-wise polymerization score is derived from pairwise comparisons of same-layer coaxial images, grounded in the psychophysics of relative judgment, and assembled into a Layer-wise Image Trajectory (LIT) for each print. Applied to both acellular and cell-laden formulations, LIT curves separate cleanly by post-processing outcome without any outcome label used in training, showing that fabrication quality can be predicted early in the build. Building on this signal, individual LIT curves are compared against statistical control limits derived from confirmed successful prints, enabling early detection of anomalous fabrication behavior at early-to-mid layers, well before development. To the best of the authors’ knowledge, this is the first application of in situ quality prediction and anomaly detection to cell-laden two-photon polymerization.

## 1 Introduction

Two-photon polymerization (2PP) stands out as the most advanced fabrication technology for engineering three-dimensional tissue microenvironments, being uniquely capable of producing fully freeform, cell-scale architectures with micron accuracy [1]. Its ability to spatially control material properties at the scale of individual cells makes it a powerful platform for investigating cell-cell and cell-matrix interactions, and for creating tissue models that overcome the well-documented limitations of conventional two-dimensional culture systems [2–4]. Water-based biomaterials are the benchmark choice for cell-laden biofabrication due to their tunable mechanical behavior and intrinsic ability to replicate key properties of the natural extracellular matrix (ECM) [5]. When combined with living cells, hydrogels provide a bio-compatible and supportive matrix that sustains essential cellular functions such as viability, proliferation, migration, and differentiation [5].

Nevertheless, hydrogel-based 2PP fabrication remains far less predictable than printing with conventional industrial photoresists. Identical nominal process parameters can yield substantially different outcomes across replicates, and this unpredictability is further amplified when living cells are embedded within the hydrogel during fabrication. Cells can scatter or absorb portions of the laser light, locally perturbing the polymerization kinetics in ways that are not fully controllable. Because cells are randomly distributed within the bioink, their interaction with the laser is spatially stochastic and construct-specific, meaning that each print experiences a unique local energy landscape that cannot be predicted or controlled at the parameter level. During post-processing — the development step in which unpolymerized resin is washed away to reveal the printed structure — phenomena such as swelling or shrinkage can significantly distort the printed architecture, leading to measurable discrepancies between the designed and realized geometry, often accompanied by variations in mechanical compliance that alter the functional properties of the construct [6,7]. Collectively, these factors make reliable fabrication difficult to achieve without extensive trial-and-error and limit the scalability and reproducibility of cell-laden 2PP constructs.

In situ monitoring has emerged in bioprinting literature as a strategy to evaluate construct quality by analyzing data acquired during fabrication [8–11]. Its application to 2PP, however, remains comparatively underexplored. Existing approaches have demonstrated that image data collected during the print carries information relevant to the final outcome, with classification models proposed to assign discrete quality labels to in situ image sequences or individual frames [12–15], and image-analysis methods explored to delineate polymerized regions and compare them geometrically to the expected outcome [16]. While these contributions represent important steps forward, two fundamental limitations persist. First, quality is evaluated during fabrication in isolation, without establishing what the observed state implies for the developed construct. Second, discrete labels conceal the degree and progression of polymerization: a print labelled as cured at a given layer may have accumulated very different crosslinking histories, and it is precisely this progression throughout the build that determines whether the final structure will be dimensionally stable after development.

To address these limitations, an in situ monitoring framework is introduced that tracks the continuous evolution of the fabrication state throughout the build by extracting a layer-wise polymerization score from coaxial optical images acquired during printing, with application to both acellular and cell-laden hydrogel formulations. The score is derived through pairwise image comparisons, a labelling strategy grounded in human preference theory that yields more consistent and less subjective annotations than absolute rating, and assembled into a Layer-wise Image Trajectory (LIT) that reflects what occurred during each individual print. These LIT curves are shown to cluster according to post-processing outcome, enabling both retrospective quality prediction and prospective layer-wise anomaly detection against a reference built from confirmed successful prints.

## 2 Materials and Methods

### 2.1 Ink Preparation

Two ink formulations were developed, both based on the same core composition; however, one was supplemented with a cell-loading component. Specifically, PhotoGel95% (#5208, Advanced Biomatrix) was dissolved in PBS (P4417, Sigma-Aldrich) at a concentration of 15% w/v at 40 *^◦^*C under constant stirring. LAP (900889, Sigma-Aldrich) was subsequently added as a photoinitiator at a final concentration of 0.50% wt.

For the cell-laden bioink, C2C12 cells were added to the ink formulation at 3×10^6^ cells/mL. After preparation, the resin was loaded in liquid state within a 50 mm diameter Petri dish and placed inside the bioprinting chamber. The chamber was set to a temperature of 37 *^◦^*C and a relative humidity of 95%. The substrate was loaded in such a way as to seal the bio-printing chamber. The contact between the resin and the objective, as well as the autofocus adjustment, were performed only after a waiting period of 15 minutes, allowing thermal and humidity conditions to reach stable equilibrium.

### 2.2 Cell Culture

C2C12 murine myoblasts were cultured in DMEM (D5796, Sigma-Aldrich), supplemented with 10% FBS (F7524, Sigma-Aldrich), 1% penicillin/streptomycin (A5955, Sigma-Aldrich) and 1% glutamine (G7513, Sigma-Aldrich). Cells were incubated at 37 *^◦^*C in a humidified atmosphere containing 5% CO_2_ and split upon reaching 80% confluency.

### 2.3 Two-Photon Polymerization Experiments

Two-photon laser printing was performed with a commercially available set-up (Quantum X shape, Nanoscribe GmbH & Co. KG) with a femtosecond laser wavelength of 780 nm. Printing was conducted using a 25× and 0.8 Numerical Aperture (NA) objective in a water immersion configuration.

A 130 × 130 × 50 *µ*m grid structure with 30 *µ*m square pores was designed in Autodesk Inventor, exported as an STL file, and processed using Describe software (Nanoscribe). Slicing and hatching distances were set to 0.5 *µ*m and 0.2 *µ*m, following the manufacturer’s recommendations for water-based photoresins.

A full-factorial design of experiments (DOE) was carried out to explore a broad range of printing conditions. Laser power was varied in 10 mW increments from 110 to 140 mW, and scan speed was adjusted in 20,000 *µ*m/s steps from 10,000 to 70,000 *µ*m/s. Each condition was printed in triplicate, resulting in a total of 48 prints, and all experimental runs were randomized to minimize systematic bias. Prints were performed sequentially, with each construct fabricated independently in a single continuous run. The entire experiment was conducted both in presence and absence of cells.

### 2.4 In Situ Process Sensing

During printing, in situ images were acquired upon completion of each layer, 100 per print, using the microscopic imaging system integrated into the machine. The imaging sensor is aligned coaxially with the laser objective lens, providing a magnified live view of the printing scene from above. With the ×25 magnification employed during the experiments, the system achieved a spatial resolution of approximately 0.568 *µ*m/pixel.

Despite the high nominal resolution, the acquired images exhibited artifacts that limited interpretability. The primary challenge was the low intrinsic contrast between polymerized and unpolymerized resin, which evolved during fabrication. Although contrast increased as the structure grew, accumulated out-of-focus signal from previously printed layers counterbalanced this improvement, leaving boundaries between polymerized and unpolymerized regions only partially distinguishable. In cell-laden resins, these limitations persisted, with additional scattering and occlusion introduced by suspended cells and cellular aggregates.

Despite these challenges, the overall image appearance, including regional brightness, texture, and contrast distribution, evolves in a manner that reflects the progressive accumulation of crosslinked material throughout the build, even when precise boundary delineation is not possible. The dynamic evolution of the printed structures was nevertheless observable: some prints remained largely unpolymerized throughout, others exhibited gradual polymerization, some developed features rapidly, and a subset showed bubble formation. These observations confirm that polymerization progressed at markedly different rates across process parameters, and that the image of a construct at any given layer was only meaningful in the context of the full sequence. An example of this observation is illustrated in Figure 1(b): Layer 100 of Print 2 visually resembles Layer 25 of Print 7, except for differences attributable to layer-dependent imaging effects. Viewed without layer context, the two images could be judged as reflecting equivalent polymerization states, yet they represent fundamentally different conditions: the first represents the last layer of an adequately polymerized construct, the second the early stage of a build that will develop excessive polymerization and eventually exhibit bubble formation. These observations underscore that discrete labels are insufficient to reliably predict final outcomes, highlighting the need for an in situ monitoring strategy that captures the evolving state of the build.

**Figure 1:**
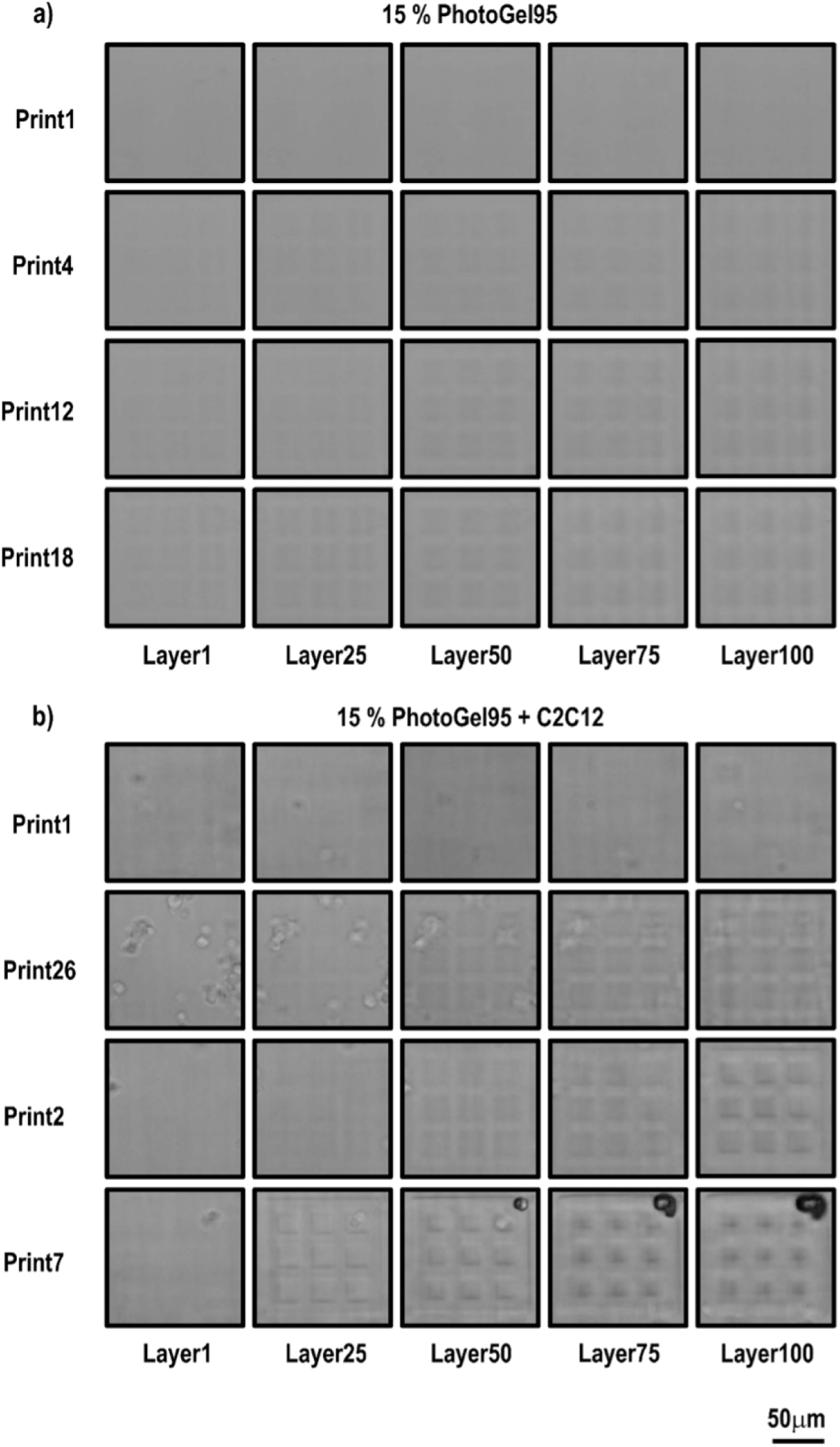
Qualitative assessment of in situ images collected during printing experiments. Images illustrate the progression of polymerization throughout layers (left to right) and across prints (top to bottom). (a) Representative images from prints using 15% PhotoGel95. (b) Representative images from prints using 15% PhotoGel95 with embedded C2C12 cells.

### 2.5 Post-Processing Evaluation

After printing, the microstructures were developed by immersing the substrates in warm PBS at 37 *^◦^*C for 5 minutes to ensure complete dissolution of the unpolymerized material. Then, each printed structure was observed using a digital imaging system (CELENA® S, Logos Biosystems) equipped with a 20× objective, achieving a spatial resolution of 0.282 *µ*m/pixel, to evaluate the integrated printing and post-processing outcome. A representative image per construct was acquired and imported in ImageJ/Fiji software for analysis. The initial evaluation was qualitative and served to identify macroscopically distinct outcomes.

As represented in Figure 2, four outcome categories were defined to summarize the range of possible post-processing results. Class 0 (absence of fabrication) corresponded to cases in which no structure was observed on the substrate, indicating either a complete lack of polymerization or polymerization that was insufficient to ensure adhesion. Class 1 (structural collapse) included samples in which partial polymerization occurred, but the intended grid geometry was not preserved, leading to irregular constructs. Class 2 (structural integrity) represented prints that retained the designed architecture without apparent defects. Finally, Class 3 (defective structures) comprised samples in which the overall geometry was maintained, but the material displayed visible signs of damage, such as burn marks or localized surface degradation. Note that, depending on the bioink type, some classes were not observed (e.g., burn marks were observed only in experiments with the cell-laden bioink).

**Figure 2:**
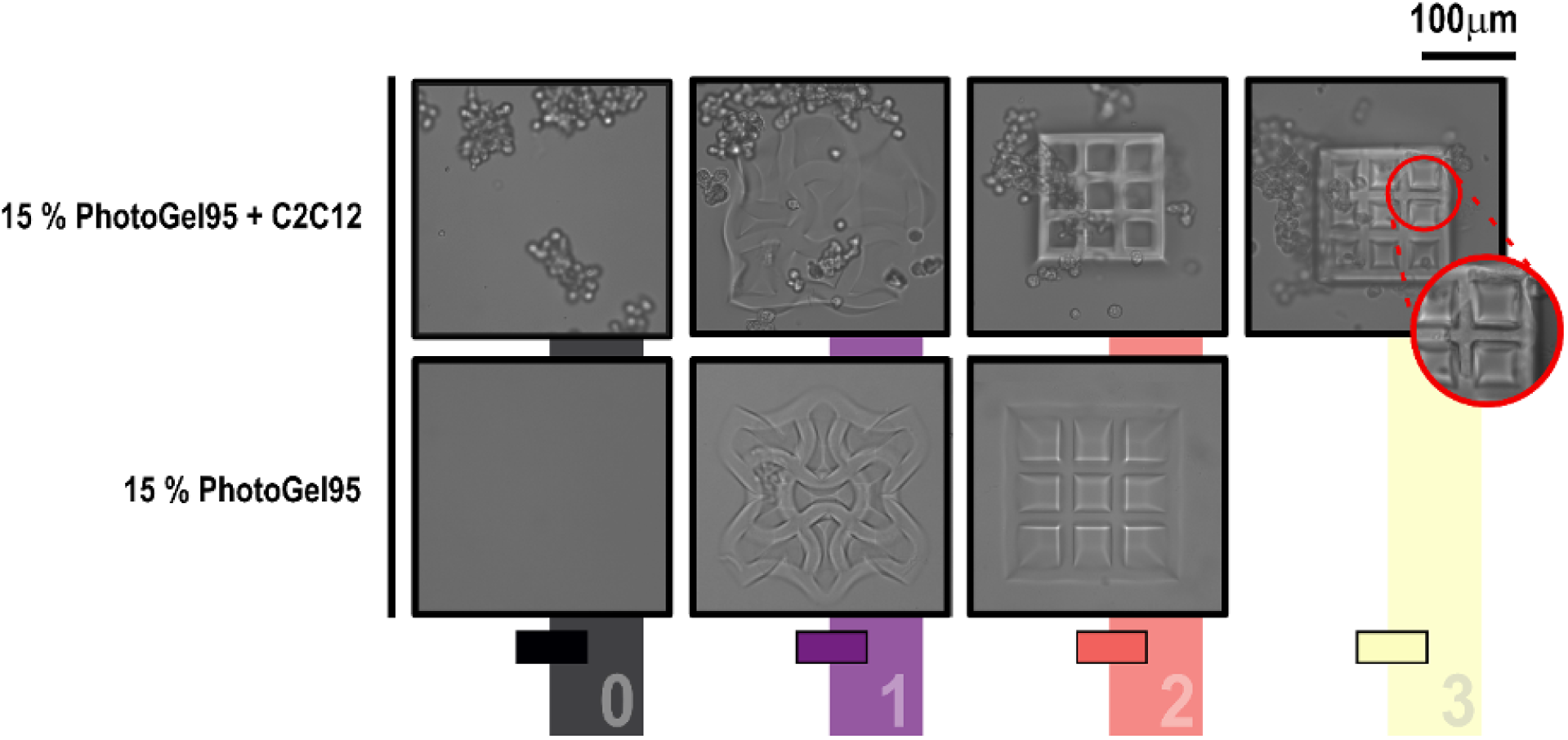
Ex situ assessment of printing results. Four different conditions were qualitatively distinguished: (black, 0) absence of a printed structure; (orange, 1) presence of a distorted grid shape; (violet, 2) detectable grid shape; and (yellow, 3) occurrence of damage during the printing process. The burn-mark defect is highlighted by a red circle; a magnified view is provided.

Quantitative image-based analysis was subsequently performed on Class 2 constructs to extract measurable parameters and provide a detailed assessment of printing performance. These constructs could still display differences in dimensions arising from swelling, an intrinsic material property that occurs independently of print quality. Since the basal layer remains anchored to the substrate, swelling at the bottom is constrained, while the upper portion expands laterally, increasing the top-view projected area without affecting the contact area at the substrate. Consequently, swelling was assessed by measuring the change in the grid area relative to the nominal area of the upper layer, as this reflects the maximal lateral expansion. For each construct, three measurements of the external area encompassing the entire construct and three independent measurements of randomly selected individual pores were performed. The mean of the replicate measurements was used for subsequent analyses. These averaged values were then incorporated into the Swelling Index (SI), where the mean pore area *Ā*_pore_ was subtracted from the external grid area *Ā*_grid_ to obtain the effective area of the construct, which was subsequently normalized over the nominal area *A*_nom_:

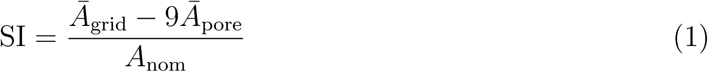

For Class 2 structures, the squareness of individual pores was calculated to provide further details and to support previous measurements. For each image, three pores were randomly selected for each condition, and for each pore, the perimeter *p* and area *A* were measured. Pore Squareness (PS) was then calculated as:

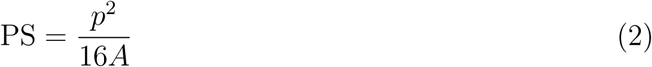

### 2.6 In Situ Data Mining of Polymerization Progression

#### 2.6.1 Model Architecture

The goal of the proposed data mining framework is to extract a meaningful signal from in situ images that reflects how far polymerization has progressed at each layer. One natural approach would be to segment polymerized from unpolymerized regions directly from the images; however, segmentation assumes that polymerization state is fully captured by spatial extent, whereas the images carry additional information in pixel intensity and local contrast that reflects crosslinking density rather than binary presence or absence of material. Beyond this conceptual limitation, the imaging artifacts described in Section 2.4 make reliable boundary delineation impractical even for a human annotator. A second approach would be to classify each image into a discrete class associated to a polymerization state; however, this would be highly subjective and strongly layer-dependent, as the same degree of polymerization manifests very differently at different stages of the build, making any absolute label assigned to a single image inherently ambiguous and difficult to define consistently.

Therefore, the problem was framed as the indirect measurement of a latent physical quantity, polymerization progression, that cannot be directly observed but whose relative magnitude can be inferred from image appearance. Direct measurement of polymerization state during fabrication is physically inaccessible, and absolute visual labelling is unreliable because the same image appearance carries different meaning at different stages of the build. Relative judgments, by contrast, are well-defined and consistent: given two images acquired at the same layer, a human expert can reliably determine which reflects more advanced polymerization based on observable image features such as contrast, boundary sharpness, and pore definition. This observation is grounded in human preference theory, which establishes that comparing two stimuli directly yields more consistent and reliable judgments than assigning absolute scores, a well-documented principle in psychophysics and perceptual evaluation [17]. In the context of in situ image annotation, where absolute polymerization state is ambiguous and layer-dependent, pairwise comparisons between same-layer images provide a more tractable and reproducible labelling task than absolute scoring.

To implement this, a Siamese architecture was adopted [18], in which two images were processed independently by a shared-weight encoder *f_θ_*, each mapped to a scalar embedding *s* = *f_θ_*(*x*) ∈ R. We employed ResNet18 [19] as *f_θ_*, yielding a 512-dimensional feature vector after global average pooling. This feature vector was then passed through a fully connected layer to produce a scalar embedding. The pair of images (*x_A_, x_B_*) was associated with a ranking label *y* ∈ {−1, 0, +1}, where *y* = +1 indicates more advanced polymerization in the first image, *y* = −1 in the second, and *y* = 0 indicated indistinguishable fabrication states. The network produced *s_A_* = *f_θ_*(*x_A_*) and *s_B_* = *f_θ_*(*x_B_*), and was trained such that ideally *s_A_ > s_B_* when *y* = +1, *s_A_ < s_B_* when *y* = −1, and *s_A_* ≈ *s_B_* when *y* = 0.

For an image pair with embeddings *s_A_* and *s_B_* and label *y*, the ranking loss was defined as a margin-based hinge loss adapted to operate on scalar embeddings and to incorporate three-way ordinal labels:

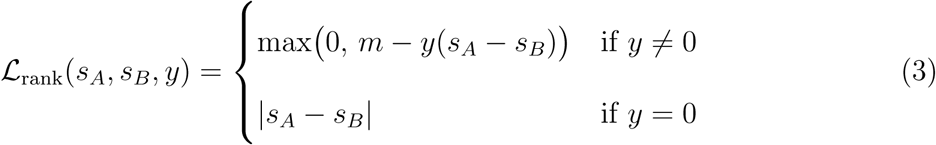

For ordered pairs (*y* ≠ 0), the loss penalized predictions where the score difference falls below a minimum margin, enforcing a minimum separation between correctly ordered embeddings and driving the encoder to identify image features that are diagnostic of polymerization differences. For tied pairs (*y* = 0), the loss reduced to the absolute difference between embeddings, directly penalizing any separation between images assigned equal fabrication state and enforcing invariance to image differences that do not reflect distinct levels of polymerization progress.

#### 2.6.2 Data Splitting and Labelling

For each bioink formulation (PhotoGel95% and PhotoGel95% with embedded C2C12), the collected dataset comprised 4800 in situ images (48 prints × 100 layers). After acquisition, each image was cropped to a 256 × 256 pixel region of interest depicting the printed area and converted to grayscale. To improve the visual clarity of the acquired images, a two-step preprocessing pipeline was applied: median blur filtering to suppress high-frequency noise, followed by Contrast Limited Adaptive Histogram Equalization (CLAHE) to locally enhance contrast while preventing over-amplification in low-signal regions.

For each bioink condition, the images were partitioned at the print level into a held-out test set, comprising 12 prints (25%), and a training pool, comprising the remaining 36 prints (75%). The training pool was further split into 5 cross-validation folds (7, 7, 7, 7, and 8 prints). Given that with only 48 prints purely random splitting could produce partitions where certain process outcomes were entirely absent, the qualitative post-processing classes identified in Section 2.5 were exploited for stratification during splitting. Importantly, the post-processing class was never used as a model input or supervision signal of any kind.

Following the print-level split, pairwise ranking samples were generated independently within each partition (Figure 3). For each layer, unique pairs were sampled without replacement, always combining images from two different prints at the same layer index. Cross-layer comparisons were explicitly excluded as they would conflate inter-print quality differences with the natural temporal evolution of polymerization. To maintain the 75/25 split at the pair level as well, 5 pairs per layer were selected from the held-out test set and 3 from each of the 5 training folds (15 in total). Across 100 layers, this yielded 500 pairs in the held-out test set and 1500 in the cross-validation pool.

**Figure 3:**
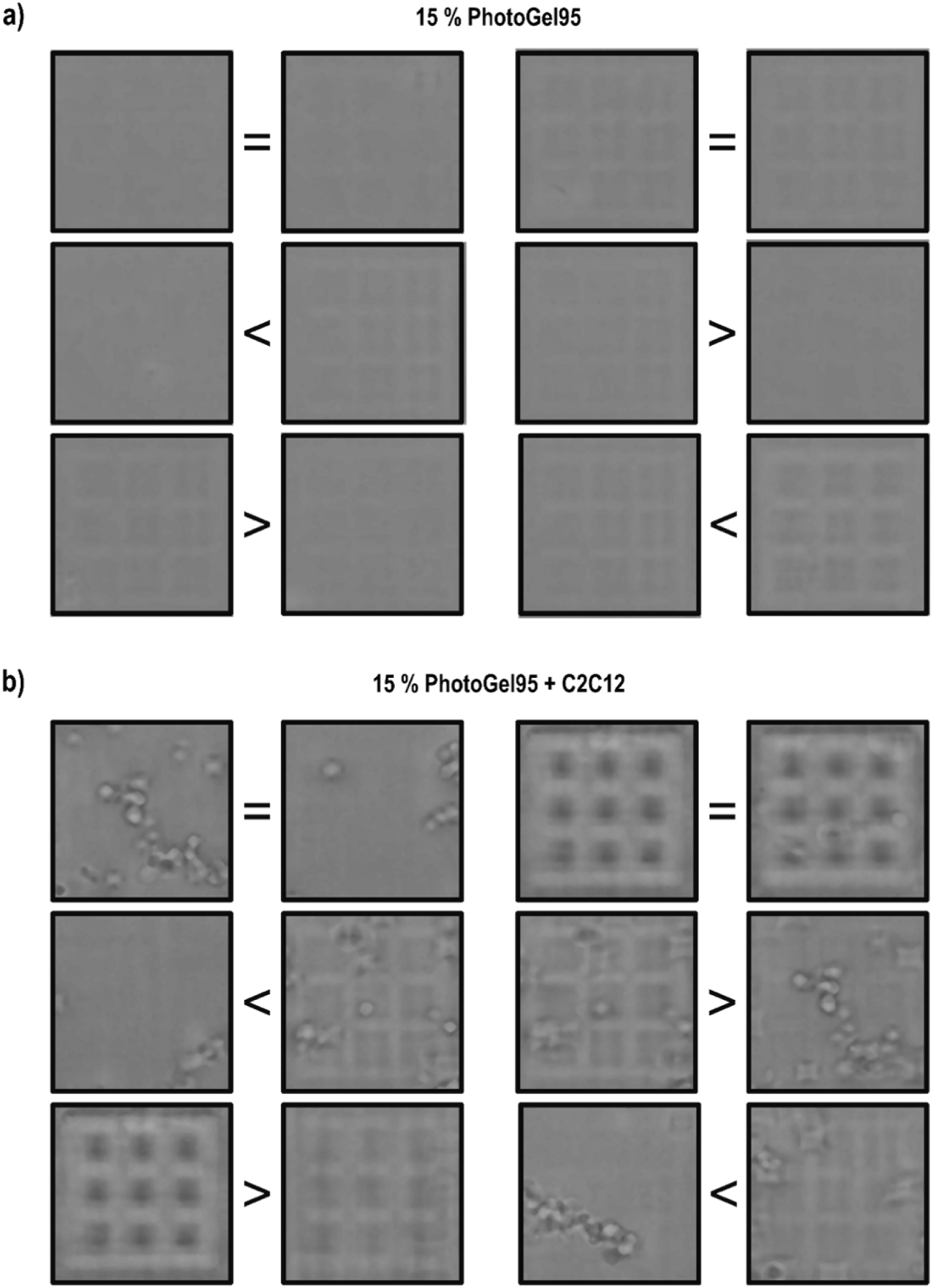
Representative image pairs and corresponding labels from the dataset. Symbols (*>*, *<*, =) indicate the relative degree of polymerization between each image pair. (a) Example pairs from samples printed with 15% PhotoGel95. (b) Example pairs from samples printed with 15% PhotoGel95 containing embedded C2C12 cells.

The ranking label assigned to each pair was determined solely by direct visual comparison of the two same-layer images, guided by three observable criteria. The first was the overall sharpness and contrast of the polymerized regions relative to the surrounding unpolymerized resin, where lower contrast and blurrier boundaries were interpreted as indicative of lesser fabrication progress. The second was pore boundary definition, where crisp, geometrically regular edges were associated with more advanced polymerization and diffuse or irregular boundaries with lesser progress. The third was apparent pore size, considered when the first two criteria were insufficient to establish a clear ranking, where visually smaller pores were associated with greater polymerization. Based on these criteria, if a clear difference was discernible the pair was assigned *y* = +1 if the first image ranked higher or *y* = −1 if the second ranked higher; if no clear distinction could be established the pair was treated as a tie and assigned *y* = 0.

#### 2.6.3 Model Training

During the training phase, four margin values (*m* = 0.1, 0.25, 0.5, 1.0) were evaluated. Each value was assessed using 5-fold cross-validation, with training consisting of 25 epochs per fold using the Adam optimizer (learning rate 1 × 10*^−^*^4^) to minimize the ranking loss. During data loading, data augmentation was applied on-the-fly. For pairwise inputs, augmentations were applied independently to each image. The pipeline included random resized cropping (scale range: 0.8–1.0), random horizontal and vertical flipping, random rotation within ±30*^◦^*, and brightness and contrast jitter within ±20%.

Performance was evaluated using Kendall’s *τ* -b, averaged across folds. Kendall’s *τ* -b is a non-parametric rank correlation coefficient ranging from −1 (perfect disagreement) to +1 (perfect agreement), with 0 indicating chance-level performance, and includes a correction for tied pairs. In addition, the pairwise ranking accuracy was computed as the proportion of correctly ordered pairs after excluding ties.

**Table 1:** Cross-validation results for 15% PhotoGel95.

| $m$ | Kendall’s τ-b | Rank Accuracy |
| --- | --- | --- |
| 0.1 | $0.648 \pm 0.088$ | $0.922 \pm 0.076$ |
| **0.25** | **$0.682 \pm 0.070$** | **$0.935 \pm 0.047$** |
| 0.5 | $0.655 \pm 0.089$ | $0.917 \pm 0.067$ |
| 1.0 | $0.621 \pm 0.129$ | $0.895 \pm 0.096$ |

**Table 2:** Cross-validation results for 15% PhotoGel95 with embedded C2C12.

| $m$ | Kendall’s τ-b | Rank Accuracy |
| --- | --- | --- |
| 0.1 | $0.686 \pm 0.079$ | $0.963 \pm 0.041$ |
| **0.25** | **$0.699 \pm 0.055$** | **$0.974 \pm 0.024$** |
| 0.5 | $0.685 \pm 0.053$ | $0.964 \pm 0.020$ |
| 1.0 | $0.685 \pm 0.074$ | $0.959 \pm 0.036$ |

Across both datasets, a margin of *m* = 0.25 yielded the best overall performance, achieving the highest Kendall’s *τ* -b and rank accuracy in cross-validation. The final model was retrained on the full training pool using *m* = 0.25 and evaluated on the held-out test set. On the 15% PhotoGel95 dataset, the model achieved a Kendall’s *τ* -b of 0.769 and a rank accuracy of 0.992. On the dataset containing embedded C2C12 cells, the model achieved a Kendall’s *τ* -b of 0.754 and a rank accuracy of 0.963. These results are consistent with strong performance on unseen samples while reflecting the benefits of training on the full dataset.

#### 2.6.4 Polymerization Score Evolution Curves

For each bioink formulation, the final model was employed to process the complete layer sequence of each print, with the encoder mapping each of the 100 in situ images to a predicted polymerization score and producing a time series reflecting the progressive evolution of the fabrication state as layers were built. Due to inherent noise, the scores were smoothed using an Exponential Moving Average (EMA):

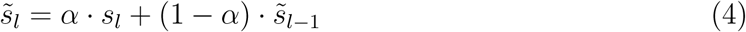

where *s_l_* is the raw score at layer *l*, *s̃_l_* is the smoothed score, and *α* = 0.3 is the smoothing factor. EMA was selected over batch smoothing methods because the smoothed value at any layer depends exclusively on current and past observations, making it directly compatible with layer-by-layer real-time deployment without requiring the full sequence to be available in advance. The resulting smoothed time series constitutes the Layer-wise Image Trajectory (LIT) for each print.

#### 2.6.5 In Situ Process Monitoring Demonstration

To illustrate the monitoring potential of the framework, a reference interval was constructed from the LIT curves of all prints with confirmed structural integrity, defined as constructs that retained the designed architecture after development, irrespective of swelling extent. The pointwise mean *µ_l_* and standard deviation *σ_l_* of the EMA-smoothed scores were computed across these reference curves at each layer *l*, defining layer-wise upper and lower control limits (UCL and LCL, respectively):

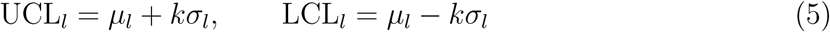

where *k* was set to control the familywise false alarm rate at 5% across all 100 monitored layers, accounting for simultaneous comparisons via Bonferroni correction. The EMA-smoothed LIT of each monitored print was then compared against this band layer by layer, with the first layer at which the smoothed score exceeded UCL*_l_* or fell below LCL*_l_* recorded as the alarm layer. Prints whose curves remained within the band throughout the full sequence were classified as non-alarming. For each bioink formulation, a representative set of prints was selected to illustrate out-of-control behavior.

## 3 Results

### 3.1 Two-Photon Polymerization Post-Processing Outcomes

Figure 4 summarizes the qualitative evaluation of the post-processed prints across both bioink formulations. Distinct behaviors were observed between conditions. The acellular Photo-Gel95% formulation yielded predominantly Class 0 outcomes across the parameter space, with structural integrity achieved only at the lowest tested scan speed (10,000 *µ*m/s) and higher laser powers (130–140 mW). In contrast, the cell-laden formulation displayed a substantially broader range of conditions enabling successful construct formation, including Class 2 and Class 3 outcomes at lower energy doses, consistent with cells effectively increasing the local energy dose delivered to the bioink during fabrication. Across the 48 prints per formulation, the cell-laden condition yielded 23 Class 0, 9 Class 1, 10 Class 2 and 6 Class 3 constructs, while the acellular condition yielded 36 Class 0, 7 Class 1 and 5 Class 2 constructs, with no Class 3 observed.

**Figure 4:**
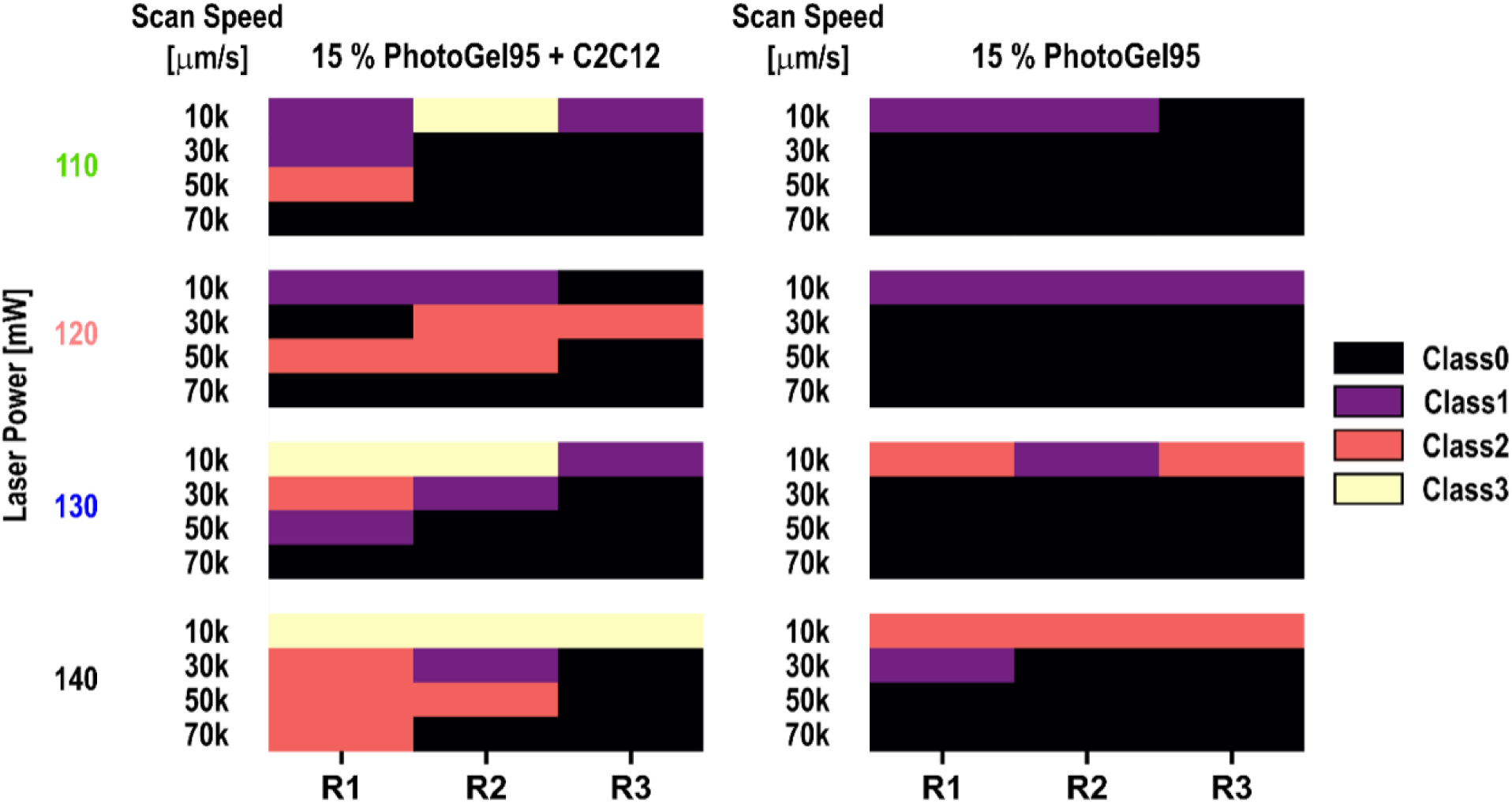
Summary table of the observed conditions under the different printing parameters tested. For each laser power and scan speed combination the results for the three replicates (R1, R2 and R3) are shown. Results obtained with PhotoGel95 laden with C2C12 cells are shown on the left, while results obtained with cell-free PhotoGel95 are reported on the right.

Despite these overall trends, the most striking observation is the variability within individual parameter combinations. As evident from Figure 4, multiple parameter settings yielded all three or even four distinct outcome classes across the three replicates, meaning that nominally identical prints produced outcomes ranging from complete fabrication failure to structural integrity or damage within the same experimental condition. This level of construct-level unpredictability cannot be attributed to parameter choice and cannot be resolved through parameter refinement alone. It directly confirms that what determines the outcome is not what parameters were set, but what occurred during each individual print, establishing the core motivation for construct-level in situ monitoring.

Figure 5 reports the swelling index (SI) and pore squareness (PS) for constructs that retained structural integrity. Cell-laden constructs generally exhibited SI values closer to the nominal reference, indicating better dimensional fidelity, whereas acellular constructs displayed systematic over-swelling. Pore squareness followed the same trend: acellular constructs showed greater and more irregular pore deformation, while cell-laden constructs presented mild but consistent deviations from ideal square geometry, suggestive of slight over-polymerization.

**Figure 5:**
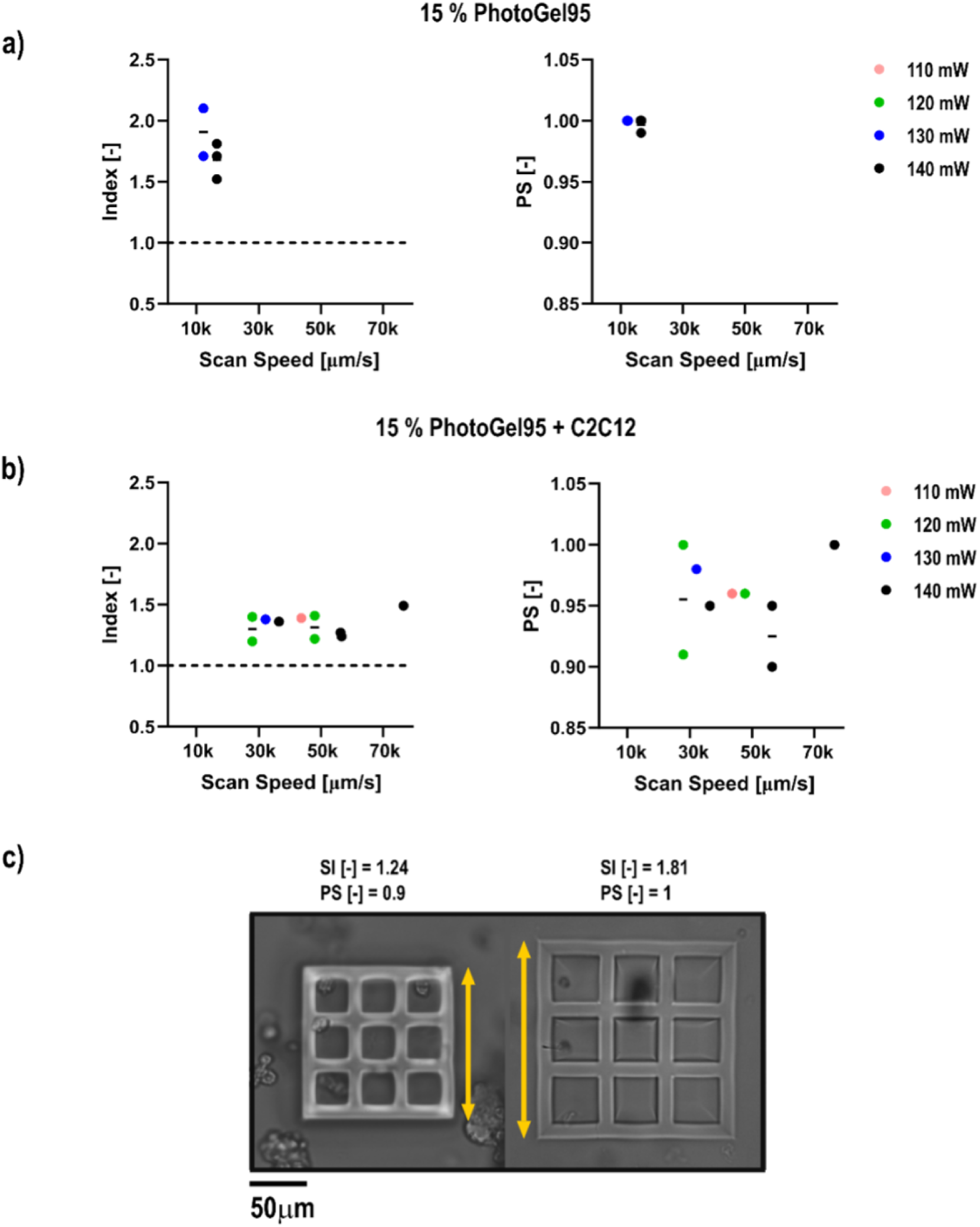
Swelling index (SI) and pore squareness (PS) values measured under Class 2 conditions. (a) Results for cell-laden samples; (b) Results for cell-free samples. The black bar indicates the mean value. (c) Visual representation of two samples with related SI and PS. Yellow arrows indicate the observed swelling, highlighting changes in structure size between samples.

### 3.2 LIT Curves

Figure 6 shows the mean LIT curves and associated ±1*σ* interval computed across all prints within each outcome class, for both bioink formulations. Despite the inherent imaging challenges described in Section 2.4, the EMA-smoothed curves reveal clear and consistent differences in fabrication behavior across outcome classes. These differences emerge directly from what occurred during each individual print, independently of the nominal process parameters used.

**Figure 6:**
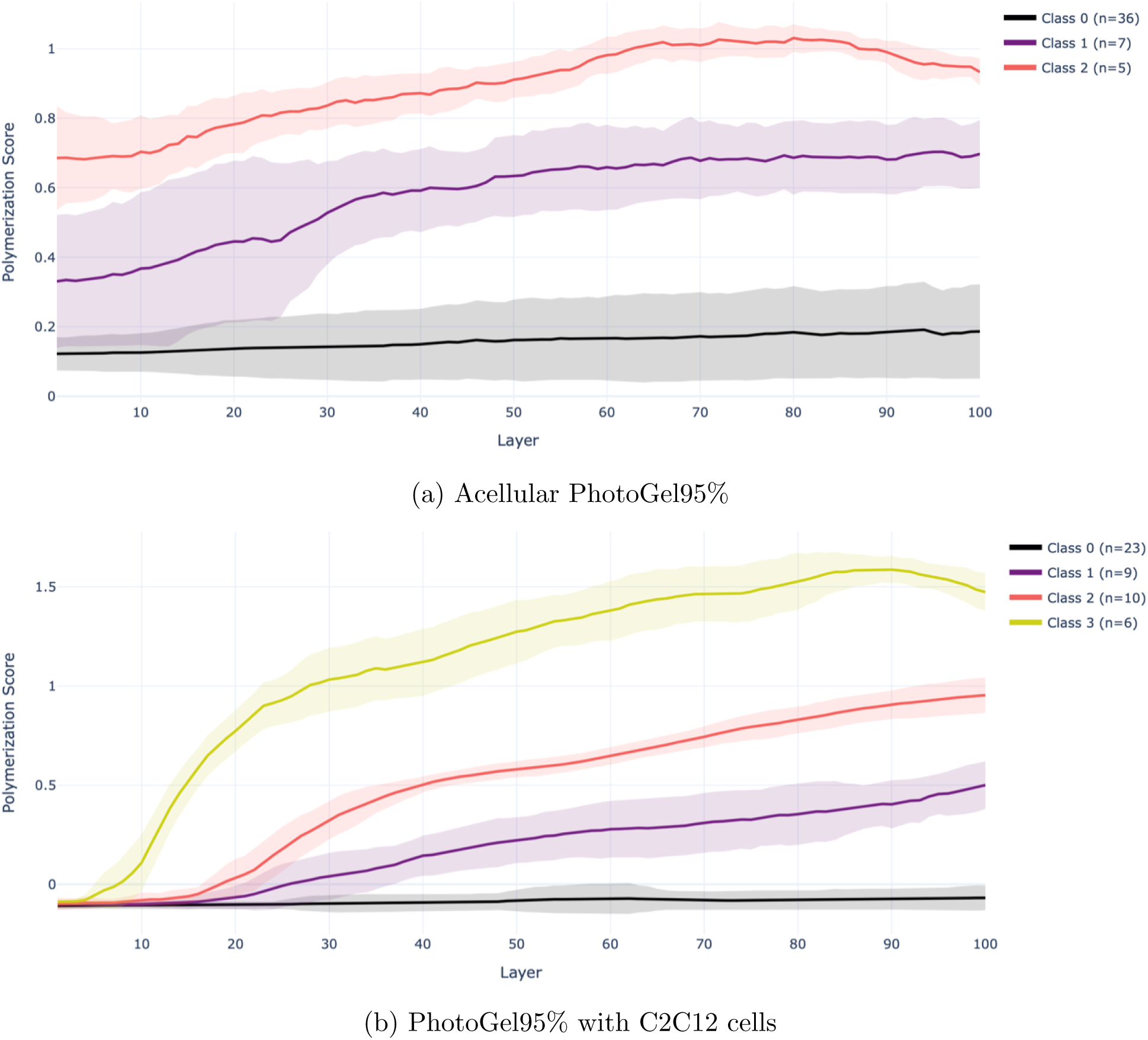
Mean LIT curves and associated ±1*σ* intervals computed across all prints within each outcome class. Curves are smoothed using a causal Exponential Moving Average (*α* = 0.3). Solid lines indicate the class mean at each layer; shaded bands indicate ±1*σ* across prints within the class. Class counts are reported in the legend. Colors correspond to outcome classes: Class 0 — absence of fabrication; Class 1 — structural collapse; Class 2 — structural integrity; Class 3 — defective structures.

In the acellular condition (Figure 6a), class separation is apparent from the earliest layers and persists throughout the build. Class 2 prints maintain clearly elevated scores from the start, reflecting homogeneous crosslinking throughout the build. Class 0 prints, which represent the large majority of the acellular dataset, remain low and nearly flat throughout, indicating negligible polymerization progression. Class 1 prints occupy an intermediate range, consistent with partial polymerization sufficient for substrate adhesion but insufficient to preserve the intended geometry.

In the cell-laden condition (Figure 6b), class separation is more pronounced and develops earlier. Class 3 prints exhibit a steep upward trajectory from the earliest layers, reaching scores substantially above all other classes by layer 20, consistent with rapid and excessive polymerization leading to visible damage. Class 2 prints show a sustained and progressive increase throughout the build. Class 1 prints rise more slowly and plateau at intermediate values, consistent with partial polymerization sufficient for substrate adhesion but insufficient to preserve the intended geometry. Class 0 prints remain effectively flat throughout, indicating negligible polymerization progression from start to finish.

Across both conditions, the score ordering follows the expected physical progression from insufficient to excessive polymerization, confirming that the framework extracts a meaningful and ordered signal of fabrication state. Critically, this ordering emerges without any class information having been used during model training, meaning that the signal is grounded entirely in image-observable features of polymerization progression, and its correlation with post-processing outcomes is an emergent property of the framework rather than a supervised result. The fact that LIT curves cluster according to post-processing outcome without any class supervision directly motivates their use as a monitoring signal, as explored in the following section.

### 3.3 In Situ Process Monitoring Demonstration

Figure 7 illustrates the monitoring potential of the framework for both bioink formulations. The UCL and LCL capture the expected evolution of the polymerization score throughout the build. The faint blue traces show the individual reference prints, and the dashed line indicates their layer-wise mean. A representative selection of additional prints is overlaid against this band, with each trace transitioning from gray to red at the alarm layer, corresponding to the first layer at which the smoothed score exits the control limits.

**Figure 7:**
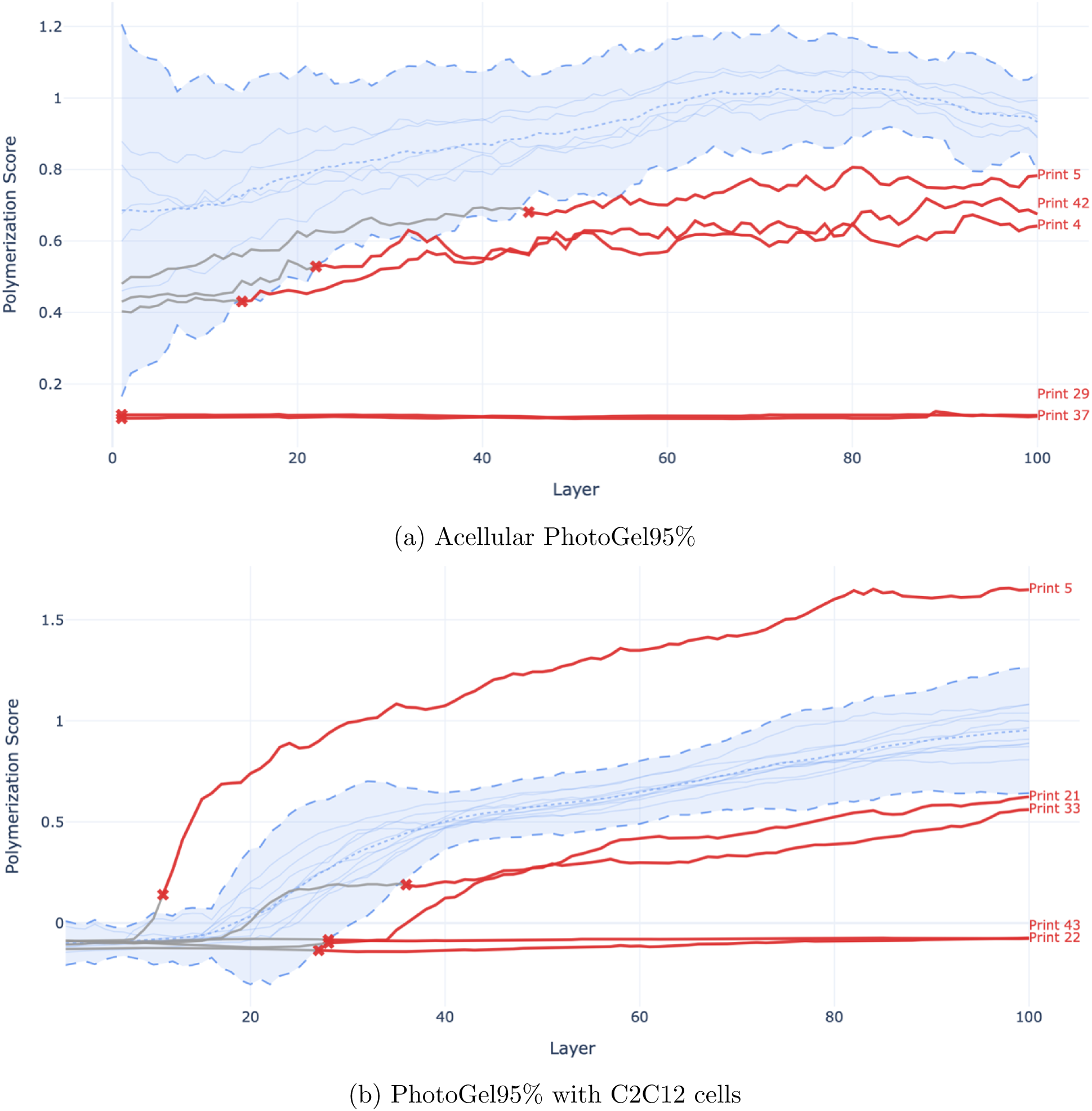
Process monitoring demonstration. The shaded region defines the reference band (mean ± *kσ*, Bonferroni corrected at familywise *α* = 0.05 across 100 layers) derived from prints with confirmed structural integrity. The dashed line indicates the layer-wise mean of the reference prints; faint blue traces show their individual EMA-smoothed LIT curves. Gray segments indicate layers within the reference band; red segments indicate layers following the alarm point, marked by an × symbol. All curves are smoothed using a causal Exponential Moving Average (*α* = 0.3).

In the acellular condition (Figure 7a), Prints 29 and 37 are flagged from the very first layer, their scores falling entirely below the reference band throughout the build, consistent with the near-complete absence of polymerization characteristic of Class 0 in this formulation.

Prints 4 and 42 exit above the band in early layers, while Print 5 exits below at layer 44. The reference band is notably wide in the earliest layers, reflecting the greater variability among reference prints at the start of the build, before tightening as polymerization stabilizes. In the cell-laden condition (Figure 7b), deviations are detected early and in both directions. Print 5 exits the upper control limit at layer 11, consistent with excessive polymerization leading to visible damage. Prints 43, 22 and 33 exit below the lower control limit before layer 30, reflecting insufficient polymerization and structural failure. Print 21 also exits above the band but later and less abruptly, suggesting a milder degree of over-polymerization.

These results demonstrate that monitoring individual print trajectories against a reference built from confirmed successful prints enables early detection of anomalous fabrication behavior, providing an actionable signal at a point when intervention is still possible.

## 4 Discussion

The ex situ evaluation revealed clear differences in post-processing behavior between the two bioink formulations. The cell-laden formulation yielded constructs with confirmed structural integrity across a broader range of printing conditions and at lower nominal energy doses, while the acellular formulation required higher and narrower parameter combinations to achieve comparable results. In terms of dimensional fidelity, cell-laden constructs exhibited swelling index values closer to the nominal reference, whereas acellular constructs displayed systematic over-swelling and greater pore deformation. These observations point to a common underlying cause: the presence of cells effectively increases the local energy dose delivered to the bioink during fabrication, promoting more extensive crosslinking under equivalent nominal parameters. While this shifts the processable window towards lower energy doses, it also introduces a higher risk of over-polymerization at the upper end of the parameter space. These differences in polymerization behavior between formulations represent a finding in their own right, confirming that the presence of cells fundamentally alters the fabrication process and that acellular and cell-laden conditions cannot be treated as equivalent, each requiring independent characterization and monitoring.

Critically, the most important finding of the ex situ evaluation is not the difference between formulations but the variability within them. As shown in Figure 4, identical parameter combinations simultaneously yielded qualitatively distinct outcomes within the same replicate set, including cases where structural integrity and complete fabrication failure were observed side by side. This variability is not attributable to parameter choice and cannot be resolved through parameter refinement. Its origin lies in the construct-specific nature of the fabrication process: in cell-laden bioinks, the spatial distribution of cells within the resin is random and uncontrollable, meaning that each print experiences a unique local energy landscape that no global parameter setting can account for. Even in the acellular case, sensitivity to local physicochemical conditions introduces a degree of stochasticity that persists regardless of nominal settings. What determines the outcome is therefore not what parameters were set, but what occurred during each individual print — and this can only be resolved through direct observation of the fabrication state.

The proposed framework addresses this directly. The Siamese model successfully extracted a continuous polymerization score from in situ images despite the imaging artifacts that preclude reliable segmentation or absolute classification. Crucially, because the model was trained exclusively on same-layer image pairs, the encoder learned to discriminate between prints at the same stage of the build without exposure to the layer-dependent appearance changes that dominate cross-layer comparisons. This design choice implicitly enforced sensitivity to inter-print polymerization differences while maintaining invariance to the natural temporal evolution of image appearance.

Two properties of this formulation jointly distinguish it from existing classification-based approaches in the 2PP monitoring literature. First, the polymerization score is continuous rather than categorical, preserving gradations in fabrication state that discrete labels collapse. Second, the framework is history-aware: the LIT encodes not only the degree of polymerization reached at any given layer but also the rate and consistency with which it evolved across the build, information that no single-layer assessment can recover. As shown in Figure 6, these properties translate into clear and physically interpretable class separation across both formulations, with the score ordering following the expected progression from insufficient to excessive polymerization without any class supervision during training.

The process monitoring demonstration in Figure 7 operationalizes these properties into an actionable signal. By comparing individual print trajectories against the UCL and LCL derived from confirmed successful prints, deviations from expected fabrication behavior are detected at early-to-mid layers, well before the build is complete and before any irreversible failure occurs. In current practice, construct quality is assessed only after development, at which point all invested resources — laser time, specialized bioink, and living biological material — are permanently lost. The framework shifts this assessment into the fabrication process itself, providing a signal that is both construct-specific and available in real time.

A subset of monitored prints exhibited trajectories that exited the reference band only partially or late in the build, suggesting the presence of a transition region where polymerization is marginal and outcomes are less predictable. This is consistent with the overlap observed between Class 1 and Class 2 in Figure 6, particularly in the acellular condition where the within-class variability of Class 1 is substantial. Expanding the reference set beyond the current dataset would help characterize this transition region more precisely and reduce uncertainty in the control limits.

## 5 Conclusions

An in situ process monitoring framework for two-photon polymerization biofabrication was introduced, capable of tracking the continuous evolution of the fabrication state throughout the build from coaxial optical images acquired layer by layer. The framework addresses a fundamental limitation of existing approaches: rather than assigning discrete quality labels at isolated points in time, it extracts a continuous polymerization score from each acquired image and assembles these into a LIT that reflects what occurred during each individual print — independently of the nominal process parameters used.

Ex situ evaluation first established that fabrication outcomes in hydrogel-based 2PP are inherently construct-specific. Identical nominal conditions simultaneously yielded qualitatively distinct outcomes within the same replicate set, a level of variability that cannot be attributed to parameter choice and cannot be resolved through parameter refinement. In cell-laden formulations this variability is further amplified by the random spatial distribution of cells within the bioink, which introduces a unique local energy landscape for each print. These findings confirm that process parameters are insufficient as quality predictors and motivate the need for a per-construct monitoring signal.

The LIT curves demonstrated clear and physically interpretable separation across outcome classes in both acellular and cell-laden formulations, with the score ordering following the expected progression from insufficient to excessive polymerization. Critically, this separation emerged without any class supervision during training, and it is an emergent property of the pairwise ranking framework grounded entirely in image-observable features of fabrication state. A process monitoring demonstration based on upper and lower control limits derived from confirmed successful prints showed that deviations from expected polymerization progression are detectable at early-to-mid layers, before the build is complete and before any irreversible failure occurs.

Two properties jointly distinguish this framework from existing classification-based approaches: the polymerization score is continuous rather than categorical, and the LIT is history-aware rather than end-point driven, together encoding information about both the degree and dynamics of polymerization that no single-layer assessment can recover. To the best of the authors’ knowledge, this represents the first application of in situ quality prediction and anomaly detection to cell-laden two-photon polymerization.

Looking ahead, expanding the dataset across a wider range of construct geometries, bioink formulations, and printing conditions remains a necessary step before the framework can be deployed as a practical process control tool. A larger and more diverse reference set would strengthen the control limits and enable more robust detection of deviations, particularly in the transition region between partial and full polymerization where outcome variability is highest. In parallel, deeper investigation into the role of cells in modifying local polymerization kinetics would inform the design of more predictable bioink systems and strengthen the physical interpretability of the monitoring signal. Ultimately, the framework is non-invasive, requires no additional instrumentation beyond the coaxial imaging system already integrated into the printer, and demonstrated consistent and interpretable behavior across both acellular and cell-laden conditions, establishing a generalizable foundation for adaptive, data-driven process monitoring in hydrogel-based two-photon polymerization biofabrication.

## Acknowledgements

This research was partially funded by the European Commission under the “HORIZON-CL4-2021-DIGITAL-EMERGING-01 project BioProS — Biointelligent Production Sensor to Measure Viral Activity” (grant agreement no. 101070120, 2022–2026).

## Notes

### Competing Interest Statement

The authors have declared no competing interest.

